# Bacterial community adaptation after freshwater and seawater coalescence

**DOI:** 10.1101/2025.09.09.675091

**Authors:** Xiu Jia, Torsten Schubert, Rick Beeloo, Aristeidis Litos, Swapnil Doijad, Pim van Helvoort, Theodor Sperlea, Matthias Labrenz, Bas E. Dutilh

## Abstract

Microbial community coalescence, the merging of entire microbial communities, is common across ecosystems, particularly in estuaries where freshwater and seawater mix. The complexity of these habitats makes the *in situ* study of community dynamics after coalescence challenging, highlighting the need for controlled experiments to unravel the factors influencing the coalescence in the estuary. To study these processes, we combined natural freshwater and seawater bacterial communities at five different mixing ratios and incubated them in parallel microcosms containing freshwater or seawater incubation media. Forty mixed communities were tracked over six passages using Nanopore full-length 16S rRNA gene sequencing. In the original field samples, freshwater hosted more diverse communities than seawater. While the communities were structurally distinct, shared bacterial families accounted for approximately 95% of total reads. Many low-abundance taxa were lost upon laboratory incubation, while potentially faster-growing ones were enriched. We found that the coalescence outcome was strongly shaped by the incubation media, whereas the mixing ratio had a minor influence. Mixed communities converged toward the source community native to the incubation media, with increasing similarity at a higher source community proportion. In freshwater, a 25% inoculum of the freshwater community was sufficient to re-establish a near-native freshwater community, whereas in seawater, similarity to the native seawater community depended on the inoculation ratio. Network analysis showed a tightly connected module of seawater families, reflecting their shared habitat preference or cooperation, whereas freshwater families were more loosely connected. We also observed that most families were unaffected by mixing ratios or temporal dynamics, with only a few showing mixing ratio dependence. For instance, the freshwater family *Comamonadaceae* and seawater families *Marinomonadaceae* and *Pseudoalteromonadaceae* were dominant in their respective native environments, and increased in mixed communities in proportion to their initial source proportion. Overall, environmental filtering had a stronger impact than the mixing ratio of source communities on coalesced communities, with habitat-specific taxa further modulating the outcome. These findings advanced our understanding of microbial responses to coalescence and provided insights into microbial community assembly in dynamic estuarine systems.

**Highlights:**

- Environmental filtering outweighs the microbial source community ratio in shaping coalescence outcomes.
- Asymmetric resilience: freshwater communities require a lower inoculum to re-establish than seawater communities.
- Modularity of the seawater source community was observed during coalescence.
- Full-length 16S rRNA gene profiling with a custom dual-barcoding Nanopore protocol enables cost-effective bacterial community tracking.
- Controlled coalescence experiments offer mechanistic insights into estuarine microbial community transitions.

## Introduction

Dispersal contributes to homogenising microbial communities across habitats (Fodelianakis et al. 2019). Due to their microscopic size, microbes often disperse not as individual cells but as an entire community. When previously separated microbial communities come into contact and mix, a process known as community coalescence occurs (Rillig et al. 2015; 2016). Community coalescence has been observed in diverse systems, including flooding events, soil tillage, estuarine mixing, wastewater treatment, and microbiome transplantation, and is now increasingly recognized as a relevant concept in microbial ecology.

Once coalescence takes place, the newly formed community undergoes rapid restructuring, often leading to the emergence of an alternative community. The outcomes of this process can vary: in some cases, both source communities contribute equally, resulting in a symmetric structure; whereas in others, one dominates, leading to an asymmetric structure (Castledine et al. 2020). These outcomes are shaped by multiple factors, including selection by the new physicochemical conditions, the mixing ratio of source communities, the strength/volume of community exchange, and the temporal dynamics of coalescence events (Custer et al. 2023; Rillig et al. 2015). Moreover, post-coalescence dynamics may also be influenced by priority effects (Fukami 2015) and even newly formed species interactions (Sanz-Sáez et al. 2020), all of which modulate how communities reassemble and stabilize after merging.

Changes in salinity, pH, and nutrient availability can strongly select microbial taxa. For example, bacterial communities are highly sensitive to salinity disturbances in aquatic environments (Tammert et al. 2023). Driven by differences in growth rate and genomic adaptability, species have different niche breadths and may respond differently to environmental changes. Specialist taxa such as *Roseovarius* are uniquely adapted to high-salinity environments (Xie et al. 2025), while generalists with a broad niche breadth can persist across a wide range of habitats (von Meijenfeldt et al. 2023). For example, marine generalists can persist in freshwater and retrieve their abundance when exposed to seawater (Comte et al. 2014). Yet, the extent to which environmental selection and niche breadth influence species survival and community structure after coalescence is still insufficiently understood.

In addition to environmental filtering, the size of the source communities may influence coalescence outcomes. Larger single-species populations are more likely to establish successfully due to reduced effects of demographic stochasticity, causing propagule pressure (Acosta et al. 2015; Kinnunen et al. 2018). Likewise, a larger, diverse community also has a higher chance of establishing (Sierocinski et al. 2021). However, communities are not just the sum of independent populations. Processes such as cross-feeding or mutual facilitation among taxa can facilitate reestablishment during community assembly. Co-selection has been observed between dominant and rare species through cross-feeding during coalescence (Diaz-Colunga et al. 2022). Moreover, larger community sizes are associated with a greater number of taxa (Sierocinski et al. 2021), potentially enhancing opportunities for interspecies interactions. Nonetheless, empirical evidence for the influence of community size on coalescence outcomes remains limited.

Estuarine ecosystems provide an ideal habitat to study microbial community coalescence. This habitat experiences continuous mixing of freshwater and seawater communities along salinity gradients. Riverine inputs transport eutrophic, freshwater-associated microbiomes into the coastal sea, while tidal intrusions carry oligotrophic, marine-associated microbiomes upstream. These microbiomes are typically compositionally and phylogenetically distinct (Logares et al. 2009), and their mixing creates transition zones where both environmental conditions and microbial communities shift (Herlemann et al. 2011; Livingston et al. 2013). Salinity, in particular, has been identified as a major environmental driver influencing estuarine microbial communities, along with dissolved organic matter, temperature, and pH (Crump et al. 2004; Lucas et al. 2016; Figueroa et al. 2021). For example, *Synechococcus* tends to increase with rising salinity (Chen et al. 2019). Studies in the Baltic Sea have also shown that marine species increase in abundance with rising salinity, while freshwater species decline. For example, populations of *Candidatus* ‘Pelagibacter ubique’ (SAR11) and *Synechococcus* exhibited phylogenetic shifts along the salinity gradient (Herlemann et al. 2011). Although these studies demonstrate that salinity influences community composition, the mechanisms driving community shifts during coalescence in estuaries are still not well understood. This is especially pressing as estuarine ecosystems are particularly sensitive to sea-level rise (Mansour et al. 2018).

A small but growing number of studies have begun to reveal how diversity, stability, and dominance patterns emerge after microbial community coalescence. These include work on soil bacterial communities (Liu and Salles 2024; Huet et al. 2025; 2023), bacterial communities in aquaculture systems (Xu et al. 2024), and estuarine microeukaryotic communities (Vass et al. 2024). However, experimental studies that directly follow bacterial temporal responses to coalescence, and the factors that shape their assembly in estuarine systems remain limited. To address this gap, we experimentally mixed natural freshwater and seawater bacterial communities in defined proportions under controlled conditions. Using microcosm experiments, we tracked community dynamics over time to address two key questions: How do environmental selection and source community mixing ratios influence community establishment following coalescence? Which taxa survive or thrive in the newly coalesced communities, and do some respond consistently depending on their source?

## Materials and Methods

### Field sample collection

We selected bacterial communities from the estuarine environment because estuaries provide a freshwater-seawater transition gradient, and their conditions can be replicated easily in the laboratory using liquid medium. On 20 April 2023, we collected surface water samples from the Warnow Estuary (freshwater) and the adjacent Baltic Sea coast (brackish water, hereafter referred to as seawater), where salinity ranged from 0.41 to 13.96 parts per thousand. These two sampling sites correspond to site 1 and site 9 of a project investigating the spatiotemporal dynamics of microbial communities along a ∼30 km transect across the Warnow estuary and the adjacent Baltic Sea coast (Sperlea et al. 2025). At each site, we collected 5 L of surface water (approximately 0-30 cm depth) as described in this report, and kept the samples at 4°C before transporting them to the laboratory for further processing. We recorded sampling time, geographical coordinates, and *in situ* measurements of conductivity (as an indicator of salinity), temperature, and pH using a conductivity-temperature-depth sensor (CTD, see Supplementary Table S1). These environmental measurements are part of the published dataset (Sperlea et al. 2025).

### Coalescence experiment setup in microcosms

To investigate the influence of coalescence ratios and environmental conditions on bacterial community dynamics after coalescence, we established microcosms with five different mixing ratios of freshwater and seawater bacterial communities under two environmental conditions (freshwater and seawater).

The initial freshwater and seawater bacterial communities were prepared through stepwise filtration (Figure 1, left panel). First, organisms larger than 8 μm were removed from 5 L of water using a bottle-top filtration system. The bacterial fractions were then concentrated 25-fold to 200 ml using a 0.2 *μ*m modular tangential flow filtration (TFF) system (Vivaflow®, Sartorius, Göttingen). The filtrate of the 0.2 *μ*m TFF was further processed through a 30 kDa modular TFF (Vivaflow®, Sartorius, Göttingen) to remove living organisms, such as ultra-small bacteria, phages, and vesicles. The resulting organism-free freshwater and seawater were used as the culturing media in the microcosms. To ensure sufficient microbial biomass for analysis, we supplemented the filtered freshwater and seawater with a limited amount of sterile yeast extract and peptone, achieving a final concentration of 0.0008% yeast extract and 0.004% peptone (Sanz-Sáez et al. 2020).

**Figure 1.**
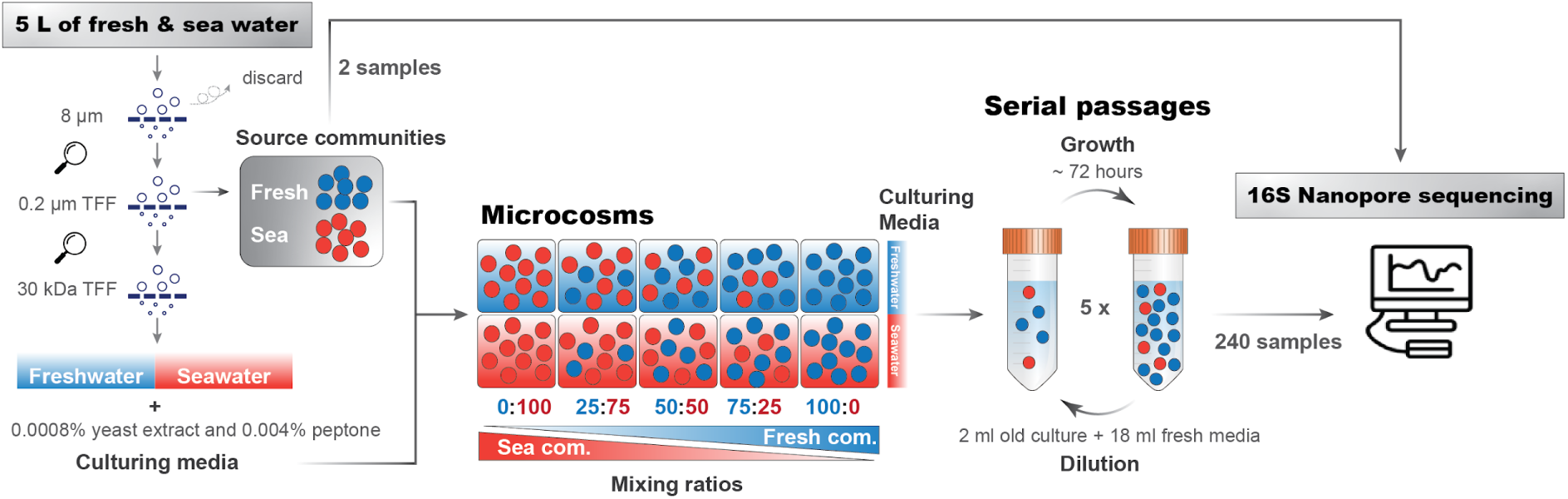
Schematic overview of the experimental setup. Microcosms were inoculated with an original or mixed community in an organism-free freshwater or seawater medium. We mixed bacterial communities from freshwater (blue dots) and seawater (red dots) sources to achieve communities with five mixing ratios. These mixed communities were cultured in either freshwater (blue background) or seawater (red background). Different combinations of the mixed microbial community and the environment are shown in squares (microcosms). Communities were transferred to fresh medium at each passage; this process was repeated five times, resulting in six passages. Changes in community composition were tracked by full-length 16S rRNA gene sequencing using Nanopore.

Community coalescence was established using five mixing ratios of bacterial communities from freshwater and seawater: 0:100, 25:75, 50:50, 75:25, 100:0 (Figure 1, center panel). To initiate community coalescence, each microcosm was prepared by adding 2 ml of either the mixed or unmixed source community to 18 ml of sterile water medium (either freshwater or seawater medium), resulting in a total volume of 20 ml. Each treatment was independently replicated four times, yielding a total of 40 microcosms. Here, 100% freshwater communities in freshwater media and 100% seawater communities in seawater media were referred to as reference controls for source communities.

Microcosms were incubated in dark conditions at 20 °C in an Eppendorf Innova 44R incubation shaker, with continuous shaking at 200 rpm under standard atmospheric pressure and oxygen levels. The incubation temperature was selected to reflect the average *in situ* summer conditions at the sampling site (Sperlea et al. 2025). Bacterial communities in our microcosms reached a relatively stable cell density within approximately 72 hours, as determined by OD600 and SYBR green measurements (Supplementary Figure S1). Accordingly, 72 hours was used as the standard interval between passages, except for passage 1, which was sampled at 92 hours after inoculation. At each transfer, 2 ml (10% of the culture volume) was inoculated into 18 ml of freshwater or seawater medium to initiate the next passage. This transfer process was repeated five times, resulting in a total of six passages. At each passage, an uninoculated medium control (n = 1) was incubated under identical conditions as a medium negative control (Supplementary Figure S2).

At the end of each passage, before the next transfer, 10 ml of each microcosm was collected to monitor community dynamics over time. Samples were centrifuged at 13,000 × g for 2 min to pellet bacterial cells. The supernatant was discarded, and bacterial pellets were stored at −80 °C for subsequent DNA extraction.

### DNA extraction and full-length 16S PCR sequencing

Bacterial pellets were resuspended in TE buffer, and cells were lysed stepwise using lysozyme, proteinase K, and the lysis solution CBO from the smart DNA prep (m) DNA extraction kit (IST Innuscreen GmbH, Berlin, Germany). DNA extraction was performed according to the manufacturer’s instructions using an automated liquid handling platform, CyBio FeliX (Analytik Jena GmbH, Jena, Germany). Following chemical lysis, AMPure XP beads (Beckman Coulter Inc., USA) were added to purify DNA. DNA from the two source communities was obtained by filtering 2 L of freshwater and seawater through 0.2 µm membranes, followed by extraction using the DNeasy PowerWater Kit (QIAGEN, Germany). DNA concentration was quantified for each sample using the DeNovix dsDNA High Sensitivity Kit (DeNovix, Wilmington, USA) and CLARIOstar^Plus^ plate reader (BMG Labtech, Ortenberg, Germany).

We used full-length 16S rRNA gene sequencing to profile bacterial community composition, as it contains more information for highly resolved phylogenetic classification. Recent studies have demonstrated that Nanopore R10.4.1 provides a rapid, efficient, and affordable method for microbial identification in complex environmental samples (Zhang et al. 2023). The bacterial full-length 16S rRNA gene was amplified using the forward primer 27F (5′-AGRGTTYGATYMTGGCTCAG-3′) and reverse primer 1492R (5′-CGGYTACCTTGTTACGACTT-3′). Both primers were tagged with 17 barcodes modified from the barcode list from the Barcoding Expansion 1–96 kit (ONT, EXP-PBC096, Oxford Nanopore Technologies, Oxford, UK). A combination of these barcoded primers enables us to process up to 289 samples (see sample list with barcodes in Supplementary Table S2).

For each 16S amplification reaction, we combined 2 µl of sample DNA (average concentration ± standard error: 2.13 ± 0.16 ng/μl) with 1.25 µl of each primer (10 μM), 0.5 µl dNTPs (10 mM), and 0.25 µl Q5 High-Fidelity DNA Polymerase (New England Biolabs) for a final volume of 25 μl. To avoid single-amplification bias, each sample was amplified in three technical replicates, and the replicates were pooled before further purification and sequencing. PCR amplification was performed using a Biometra TAdvanced thermal cycler (Analytik Jena, Jena, Germany), with the following cycling conditions: 5 min at 98 °C to activate polymerase; 10 s at 98 °C for denaturation, 30 s at 60 °C for annealing, 2 min at 72 °C for extension (5 cycles); 10 s at 98 °C for denaturation, 2.5 min at 72 °C for extension (30 cycles); and a final step of 2 min at 72 °C. Potential contamination was checked by including a no-template control on each PCR plate and verifying it with gel electrophoresis. Negative PCR controls were also sequenced and yielded only a negligible number of reads compared to the biological samples (Supplementary Figure S2).

Following PCR, amplicons were purified first using AMPure beads to remove primers, dNTPs, polymerase, and other PCR reagents. Samples were quantified using the DeNovix dsDNA High Sensitivity Kit and pooled to achieve the same molecular weight. To select our target amplicons (about 1,500 bp) and remove non-target fragments, the pooled sample was further purified using the Zymoclean Gel DNA Recovery Kit (ZYMO RESEARCH, Freiburg, Germany).

The final pooled 16S rRNA library was sequenced using the Ligation Sequencing Kit V14 (SQK-LSK114, Oxford Nanopore Technologies, Oxford, UK) on an FLO-MIN114 flow cell (R10.4.1 nanopore). ZymoBIOMICS Microbial Community (Standards with even and log distribution, ZYMO RESEARCH, Freiburg, Germany) was used as a positive control of our experiment to validate whether PCR and nanopore sequencing yield the expected distribution of a targeted community (Nicholls et al. 2019). All bacterial genera from the two ZymoBIOMICS microbial communities were successfully recovered by our sequencing. The observed community composition closely reflected the expected distribution, although some deviations were present (Supplementary Figure S3). This indicates that our sequencing run can reflect the bacterial community structure, although with some variance.

### Sequence processing and feature table cleaning

After sequencing, we obtained 14.29 M raw pod5 reads (24.03 Gbases) from 272 samples, comprising 240 microcosm samples, 18 negative controls, 6 mock communities, and 8 field samples. We used Dorado (version 0.9.1) to convert raw Nanopore signals to FASTQ files. After basecalling, 14,565,367 fastq reads were retained for further analysis with a median read length of 1,554 nt and a median read quality score of 13.9. Because our barcode design is not supported by Dorado, we developed a custom tool for demultiplexing. Flanking sequences (adapters and primers) are first identified in each read using Sassy (Beeloo and Koerkamp 2025). The sequence between the adapter and primer is then compared against all barcode sequences, and a match is assigned if a barcode is found within an edit distance of ≤ 5. Since the expected amplicon length was 1300–1500 nt and the experiment used dual-end barcoding, we required that two barcode matches occur within this distance of one another for a read to be classified. To account for concatemer formation, reads containing multiples of this pattern were split accordingly. In total, 6,391,864 reads could be demultiplexed into 272 samples. The demultiplexed fastq reads were taxonomically annotated using Kraken2 (Wood *et al*., 2019) against the SILVA 182 database (Quast *et al*., 2013), with genus level as the lowest taxonomy annotation.

All further data analyses were performed using R (version 4.3.3). First, we excluded non-bacterial reads and summarised the bacterial reads at the family level, following the evidence that bacterial community structure converges at this taxonomic rank under stable conditions (Goldford et al. 2018). Next, we removed families arbitrarily with fewer than three total reads and those present in fewer than three samples to minimise the effect of potential errors in sequencing or annotation. After these steps, we retained a total of 5,371,097 bacterial reads from 357 families across 242 samples (240 microcosm samples and two field samples), with per-sample read counts ranging from 3,591 (minimum) to 83,009 (maximum), a median of 19,580, and a mean of 22,195. The rarefaction curves of most microcosm samples reached a plateau, indicating that their diversity was sufficiently captured (Supplementary Figure S4). However, the field samples did not reach a plateau, reflecting their higher diversity compared to the microcosm samples.

### Data transformation and diversity analysis

To assess how laboratory conditions influenced source community composition, we compared the log_2_ fold-change in relative abundance between natural and laboratory conditions for both freshwater and seawater habitats. The overlap of families in different habitats was plotted by https://www.deepvenn.com/. To identify freshwater and seawater bacterial families, we performed differential abundance analysis of these families between freshwater and seawater communities under laboratory conditions using the ANOVA-Like Differential Expression tool (ALDEx2), which is based on center-log ratio (CLR) transformed data.

To compare the habitat associations of the identified freshwater and seawater families with observations in other studies, we examined the samples where these families were detected in the MGnify database using the taxonomy tables of all samples processed with their 4.1 pipeline. This complete dataset consisted of 57,645 samples from 141 environments. To reduce noise and contamination, we applied a detection threshold of 50 reads to define the presence of a family in a sample. The detection rate of each family within an environment was calculated as the percentage of samples in that environment where the family exceeded the detection threshold. Of the 35 families whose habitat preference was identified by ALDEx2, 29 had associated records in MGnify. For the remaining families that were not detected at the family level in MGnify, we searched for environment associations using the most abundant genera within those families.

To address the compositional nature of amplicon sequencing data, we applied a CLR transformation to nonzero values by dividing the read counts for each bacterial family by the geometric mean of all taxa within a sample. Community dissimilarity was then quantified by calculating the Aitchison distance–the Euclidean distance between the CLR-transformed abundance of taxa across samples.

To identify taxa that co-occurred during succession following community coalescence, we performed a network analysis using SpeSpeNet (van Eijnatten et al. 2025). The analysis used a feature table that was CLR-normalised, with random pseudo-counts applied during normalisation. Families with more than five occurrences across samples were retained. Only positive correlations with Spearman’s ρ > 0.5 were included as edges in the network. The resulting network was visualised using a Fruchterman–Reingold layout with the R package ggraph. This analysis enabled the detection of taxon clusters potentially shaped by mutual adaptation to shared abiotic conditions or synergistic interactions.

## Results and Discussion

### Occurrence and habitat preferences of freshwater and seawater bacteria

This study provides an empirical analysis of coalescence between freshwater and seawater bacterial communities under controlled conditions. The two source communities showed distinct taxonomic composition in their natural environments. In samples collected upstream of the Mühlendamm of the Warnow River (freshwater) and the adjacent Baltic Sea coast at Strand Warnemünde (seawater), we detected a total of 164 bacterial families, 69 of which were shared (Figure 2A). As expected, more unique families were detected in seawater (49) than in freshwater (46), reflecting the predominant discharge from the freshwater to seawater. Families that were present in both freshwater and seawater represented approximately 95% of the sequencing reads, although their abundances differed. Habitat-unique families were comparatively rare, contributing less than 5% of the total read abundance in each (Supplementary Figure S5). The natural freshwater community was dominated by *Comamonadaceae*, *Sporichthyaceae*, *Burkholderiaceae*, *Terrimicrobiaceae,* and *Rubritaleaceae*, which were also found in seawater (Figure 2C). At the same Warnow River site, *Sporichthyaceae*, *Burkholderiaceae*, and *Rubritaleaceae* were previously identified as freshwater taxa (Kujat et al. Under review), while the dominance of *Comamonadaceae* and *Burkholderiaceae* is consistent with earlier observations from rivers in northern Germany (Clols-Fuentes et al. 2024). Similarly, *Comamonadaceae* is common in the inner Råne River estuary, which flows into the northern Baltic Sea (Figueroa et al. 2021). The seawater community from the Baltic Sea coast was dominated by *Flavobacteriaceae*, *Rhodobacteraceae*, and SAR11 Clade I, with the first two also detected in the freshwater samples (Figure 2C). Dominance of these families was also reported by other Baltic Sea studies (Herlemann et al. 2011; Dupont et al. 2014). Together, our results corroborate previous work showing that microbial community structure shifts along estuaries, where changes are often related to salinity transitions (Paver et al. 2018; Amadei Martínez et al. 2025).

**Figure 2.**
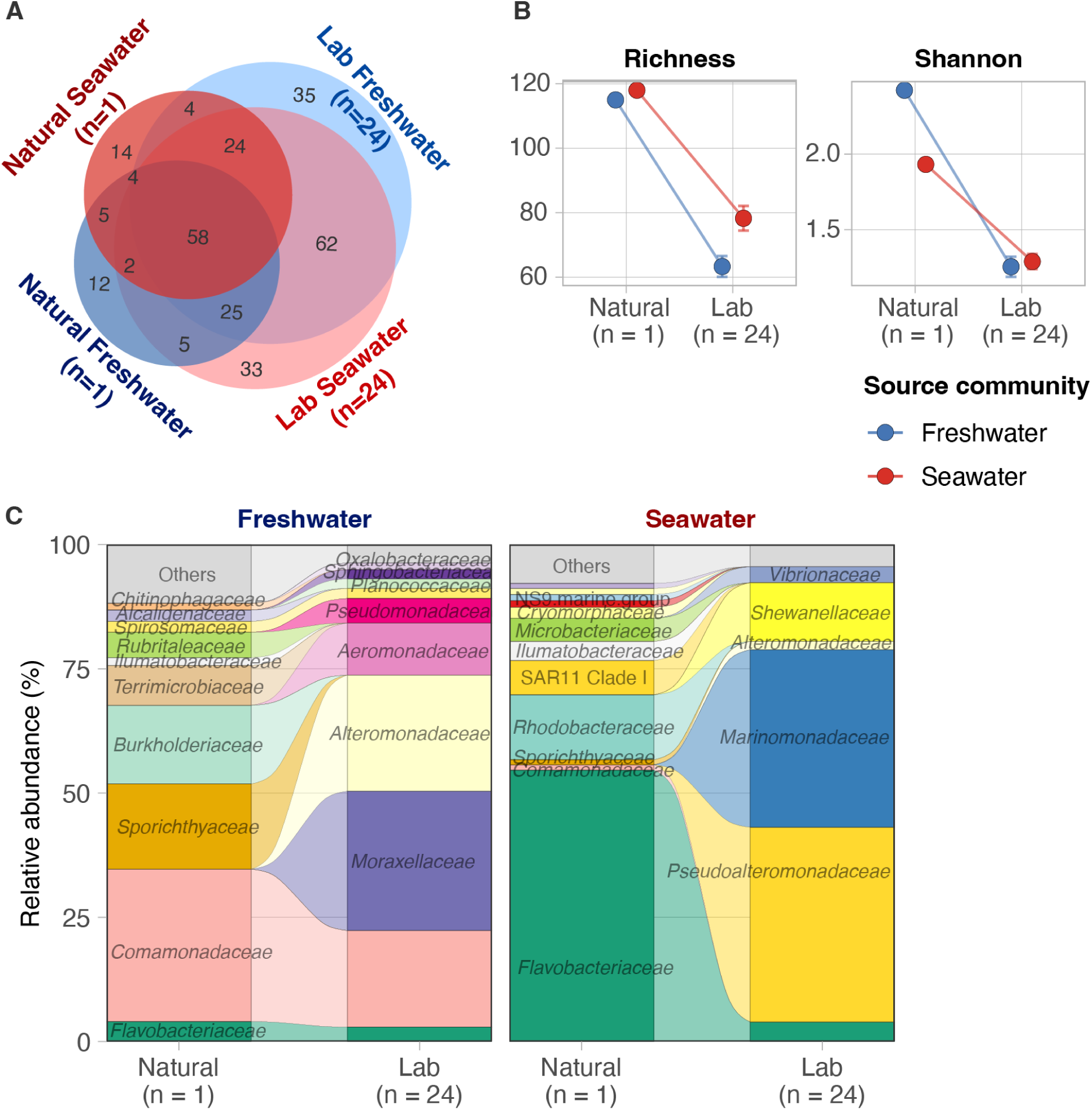
Family-level composition of bacterial communities in freshwater and seawater under natural and laboratory conditions. (A) Venn diagram showing the overlap of 294 families detected across two different water sources and two growth conditions. In natural environments, 115 families were detected in freshwater and 118 in seawater. Under laboratory conditions, 216 families were detected both in freshwater and seawater. (B) Richness and Shannon diversity of two different source communities under two growth conditions. (C) Stacked bar plots showing the relative abundance of families (relative abundance > 1%) in freshwater and seawater under natural (n = 1) and laboratory conditions (average of 24 samples).

Next, we conducted mixing experiments under controlled conditions to investigate coalescence between freshwater and seawater communities. To correct for the impact of laboratory cultivation, we also submitted the source communities to the same treatments. This allowed us to compare the dynamics of mixed and unmixed communities under similar conditions. The unmixed freshwater and seawater communities both showed reduced diversity compared to their natural counterparts (Figure 2B, Supplementary Figure S6). The total observed number is higher than the 164 bacterial families detected in the natural samples (above) because 24 samples were sequenced. Log_2_ fold change results showed that most families were depleted under laboratory conditions, with only a few becoming enriched (Supplementary Figure S7A). This reduction is likely due to several reasons. First, our 30 kDa filtration protocol removed larger particles, which can harbour higher diversity (Rieck et al. 2015). Second, by submitting the microcosms to mixing, we reduced microscale gradients, which may facilitate coexistence (Seymour et al. 2017; Schlundt et al. 2020). Third, we hypothesized that the added nutrients (0.0008% yeast extract and 0.004% peptone), although limited in amount, facilitated the outgrowth of fast growers and led to reduced diversity. Indeed, the families that were enriched after cultivation tended to have faster growth rates than those that were depleted (Supplementary Figure S7B). Together, these results show that we could cultivate distinct freshwater and seawater communities under laboratory conditions, retaining diverse bacterial families despite reduced complexity.

Under laboratory conditions, the taxonomic overlap between freshwater and seawater communities increased during six cultivation passages. Overall, 169 families were shared (64.3% of the total), accounting for 99.9% of the total sequence abundance (Figure 2A, Supplementary Figure S5). In lab-cultured freshwater communities, *Comamonadaceae, Moraxellaceae*, *Alteromonadaceae*, and *Aeromonadaceae* became dominant, whereas *Pseudoalteromonadaceae*, *Marinomonadaceae*, and *Shewanellaceae* dominated lab-cultured seawater communities (Figure 2C, Supplementary Figure S8).

To identify bacterial families with a habitat preference for freshwater or seawater under laboratory conditions, we performed ALDEx2 differential abundance analysis between freshwater and seawater reference communities. The analysis identified 16 families that were significantly enriched in laboratory freshwater (e.g., *Comamonadaceae*, *Spirosomaceae, Sphingobacteriaceae, Oxalobacteraceae, Aeromonadaceae, Alteromonadaceae, Burkholderiaceae,* and the NS11-12 marine group) and 19 families that were significantly enriched in laboratory seawater (e.g., *Marinomonadaceae, Pseudoalteromonadaceae, Shewanellaceae,* and *Vibrionaceae*; Figure 3A, Supplementary Figure S9). To compare these *in vitro* biome associations with observations in natural systems, we searched the MGnify database for the preferred environment of the enriched families (see Methods, Figure 3B). Indeed, most seawater-associated families were frequently detected in the marine environment, with exceptions including *Pasteurellaceae* and *Endozoicomonadaceae*, which were mostly detected in host-associated and mixed environments. Closer examination of the associated studies revealed that *Endozoicomonadaceae*, although identified in host-associated environments, was found on marine organisms such as seagrass (Webb et al. 2019) and sponges (Moitinho-Silva et al. 2014), confirming its preference for marine systems. Freshwater families were consistently detected in freshwater environments, but were also abundant in environments annotated as ‘Other’. While *Alteromonadaceae* and *Rubritaleaceae* are most often found in marine environments, both were more abundant in freshwater under our laboratory conditions. Although the NS11-12 marine group was not detected in the MGnify dataset, it has been identified in riverine, estuarine, and coastal systems (Amadei Martínez et al. 2025; Barberoux et al. 2025; Korlević et al. 2022).

**Figure 3.**
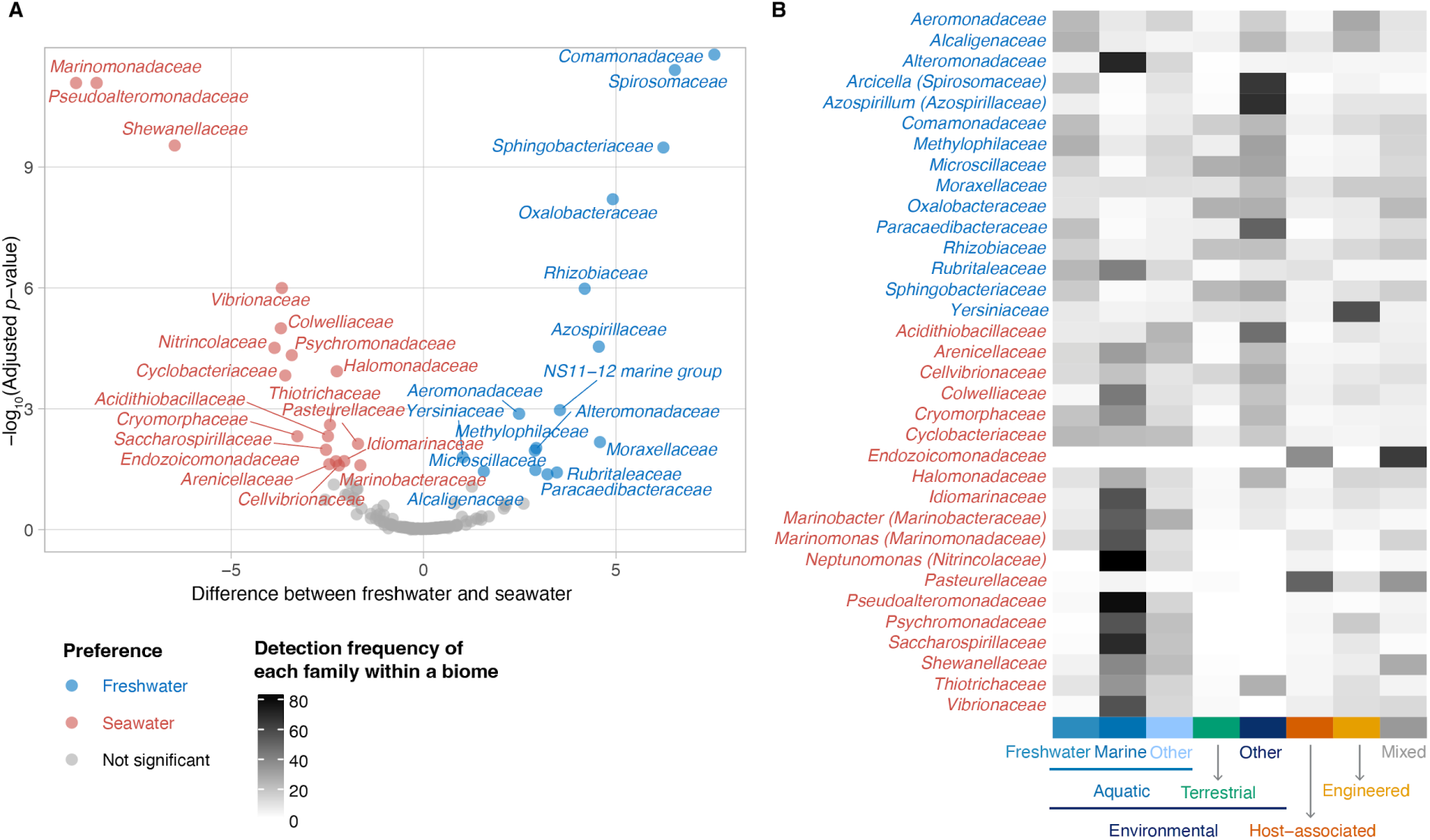
Freshwater and seawater preferred families under laboratory conditions. (A) Differential abundance of families between freshwater and seawater communities under laboratory conditions, as based on ALDEx2 analysis incorporating all 24 passages and replicates (Supplementary Data S2). The x-axis shows the median difference in CLR-transformed abundance between the two communities, and the y-axis shows the adjusted *p*-value of the Wilcoxon rank-sum test (-log_10_). Colored markers and text indicate families with significant differences (adjusted *p* < 0.05), based on the Wilcoxon rank-sum test. (B) Heatmap showing the distribution of freshwater- and seawater families across various biomes based on the MGnify database. Values represent the frequency of each family in a given biome, calculated as the number of samples in which the family was detected within a biome divided by the total number of samples in the biome.

Together, these results show that natural freshwater and seawater communities share a core set of families, yet remain compositionally distinct. While laboratory incubation reduced the diversity of both source communities, habitat associations of bacterial families after laboratory cultivation remain consistent with observations in global metagenomics data.

### Environmental conditions strongly impact community coalescence outcomes

The structure of coalesced bacterial communities was strongly influenced by the aquatic growth environments. This was evident in the clear separation of samples based on the culturing environment (Figure 4A). In freshwater, shifts in community structure were primarily driven by freshwater families such as *Comamonadaceae* and *Spirosomaceae,* while in seawater, changes were mainly influenced by seawater families like *Pseudoalteromonadaceae* and *Marinomonadaceae*. In the freshwater environment, communities containing down to 25% freshwater bacteria clustered together. Only the 100% seawater communities formed a distinct group. This was unexpected, as we anticipated that seawater communities should retain freshwater bacteria and recover. This lack of recovery may be due to threefold lower cell density in seawater (Sperlea et al. 2025), and the inoculum having few freshwater bacteria with low abundance, making reestablishment difficult. In the seawater environment, we observed a gradual shift in community structure during mixing from 100% seawater to 100% freshwater communities. These patterns suggest that in freshwater, a relatively small inoculum is sufficient to reestablish the original community and highlight either the strong influence of freshwater environments or high cell density on the structure of the coalesced community. The situation differs in seawater, where the final coalesced community reflects the mixing ratio.

**Figure 4.**
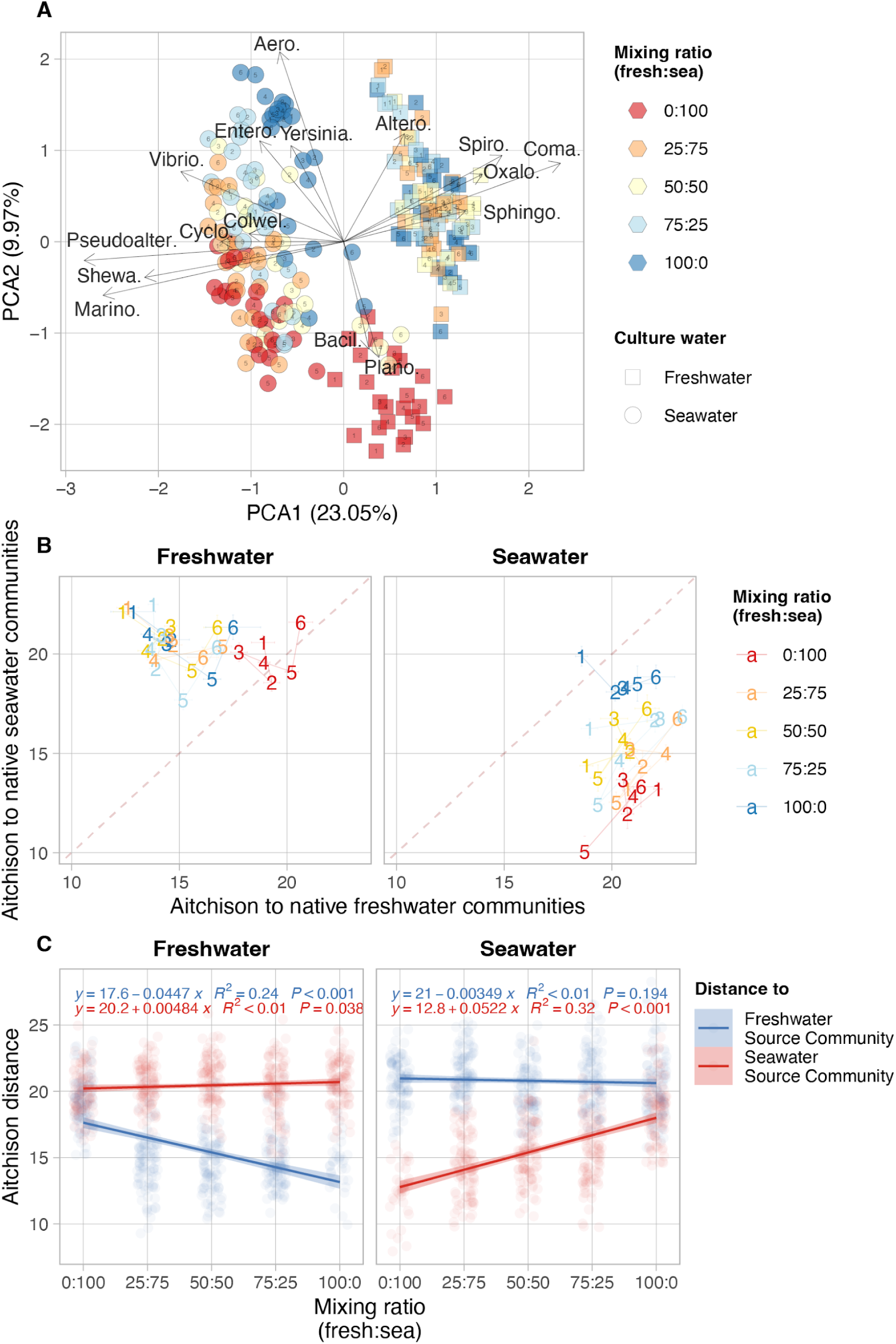
Community dissimilarity after mixing based on Aitchison distance at the family level. (A) Compositional principal component analysis (PCA) biplot based on Euclidean distance of the CLR-transformed feature table for PCA. Axes show the percentage of total variance explained by the first two components. Colors represent the mixing ratios of freshwater and seawater communities; shapes represent the growth environment; numbers inside the shapes represent passages. Grey arrows highlight families strongly contributing to variance (length > 1 on either axis), with direction and length showing the strength and associations with PCA components. Family abbreviations: Aero. – *Aeromonadaceae*, Altero. – *Alteromonadaceae*, Bacil. – *Bacillaceae*, Coma. – *Comamonadaceae*, Colwel. – *Colwelliaceae*, Cyclo. – *Cyclobacteriaceae*, Entero. – *Enterobacteriaceae*, Marino. – *Marinomonadaceae*, Oxalo. – *Oxalobacteraceae*, Plano. – *Planococcaceae*, Pseudoalter. – *Pseudoalteromonadaceae*, Shewa. – *Shewanellaceae*, Spiro. – *Spirosomaceae*, Sphingo. – *Sphingobacteriaceae*, Yersinia. – *Yersiniaceae*, Vibrio. – *Vibrionaceae*. (B) Dissimilarity (Aitchison distance) between mixed communities and their freshwater or seawater source communities. Colors represent mixing ratios; numbers represent passages. Points/Numbers represent the mean dissimilarity of the mixed community to the two source communities; error bars represent the standard error. Lines connect temporal changes across passages for each group. (C) Regression of Aichitson distance between mixed communities and their source communities as a function of the proportion of the source communities in the initial mix. Freshwater source communities refer to 100% freshwater communities cultured in freshwater; seawater source communities refer to 100% seawater communities cultured in seawater. The y-axis shows the Aitchison distance of the coalesced communities to the source communities. The x-axis indicates the initial mixing ratios of freshwater and seawater communities before culturing, with four replicates per ratio across six passages. Colors represent the source community. Left and right panels correspond to the culturing environment of mixed communities (freshwater or seawater). Regression equations, *R*² values, and *p*-values are displayed for each group in the corresponding panel.

PERMANOVA analysis confirmed that environmental conditions explained most of the community variation (*R*^2^ = 19.90, *P* < 0.001), followed by mixing ratio (*R*^2^ = 8.26, *P* < 0.001), and passages (*R*^2^ = 4.99, *P* < 0.001). Significant but weak interaction effects were also observed between mixing ratio and environment (*R*^2^ = 6.46, *P* < 0.001) and between passage and environment (*R*^2^ = 3.38, *P* < 0.001). These interactions suggest that the influence of mixing ratios and temporal changes (passages) on community structure depends on environmental context.

These findings align with previous studies and provide further evidence for the importance of environmental selection (Becking 1934). For example, mixing freshwater and seawater resulted in brackish conditions that act as a strong environmental filter (Rocca et al. 2020). In soil systems, microbes and their environment are physically linked, so they necessarily mix in equal ratios during coalescence. When soils were mixed in a recent study, abiotic factors were also found to be the main drivers of community variation. This happened as one soil imposed abiotic dominance as it had a strong buffer capacity, and disproportionally favored the establishment of pre-adapted taxa (Bresciani et al. 2025). Together, these findings support the hypothesis that when one microbial community is introduced into the environmental setting of another, environmental filtering favors bacteria that are already adapted, thereby promoting the success of resident over invading species (Rillig et al. 2015).

### Source community contributions vary with environmental conditions

To quantify the impact of the mixing ratio on the coalescence outcome, we calculated Aitchison distances of the communities to each of the source communities. In each cultivation environment, the coalesced communities were more similar to the native source community than to the introduced one (Figure 4B). When plotted against the mixing ratio, we observed that the distance to the native source communities decreased, while the distance to the introduced source communities remained relatively constant and higher (Figure 4C). This trend was especially pronounced in seawater, where the regression fit was stronger (*R*^2^ = 0.32) compared to freshwater (*R*^2^ = 0.24). These findings indicate that each environment preferentially selects for its native source community, and that increasing its proportion in the coalescence process further enhances its contribution to the coalesced community. This outcome supports the asymmetric outcome of coalescence, where one community limits the establishment of taxa from the other due to superior environmental adaptation (Castledine et al. 2020).

Similar to previous studies on community invasion dynamics, which demonstrate that higher propagule pressure enhances the chances of successful invasion (Sierocinski et al. 2021), we found that the proportion of the source community is correlated with its contribution to the coalesced community. A recent experimental and modelling study on community coalescence further supports a dose-dependent response. Interspecific competition between species from two source communities reduces the availability of shared resources, thereby driving a dose-dependent effect. The weaker competitor (i.e., the one with lower consumption rates) for these shared resources has a stronger dose-dependent response. This effect can be detected in the airwise species-mixing assay and becomes more pronounced with increasing community diversity (Goldman et al. 2025). Together, these results highlight that coalescence outcomes are shaped not only by environmental filters, but also by the initial abundance of source communities, especially when there is a strong species interaction in the source community.

### Seawater families showed stronger covariation than freshwater families during coalescence

To assess whether taxa may ecologically interact during coalescence, we constructed a co-abundance network of all bacterial families. The resulting network (modularity score = 0.225, Figure 5) revealed two main modules. The largest module, with 312 edges between 32 families, was composed of families more abundant in seawater environments, which were mainly identified as seawater families. It also showed high connectivity (average node degree = 19.5). This strong covariation not only suggests that these taxa have a similar response to environmental factors, but could also represent stronger ecological interactions. In contrast, the second module had 20 edges between 12 freshwater families, representing a module with much lower connectivity (average node degree = 3.33). The other freshwater families also formed in other small modules. This indicates a more loosely structured network among freshwater families. We did not observe strong ecological interactions between freshwater and seawater families, except for *Aeromonadaceae* and *Yersiniaceae*, which were both classified as freshwater families, but were grouped in the seawater-associated module, as they showed higher abundance when switching to another environment. This indicates that the interactions were maintained in the source communities, while new interactions were rare.

**Figure 5.**
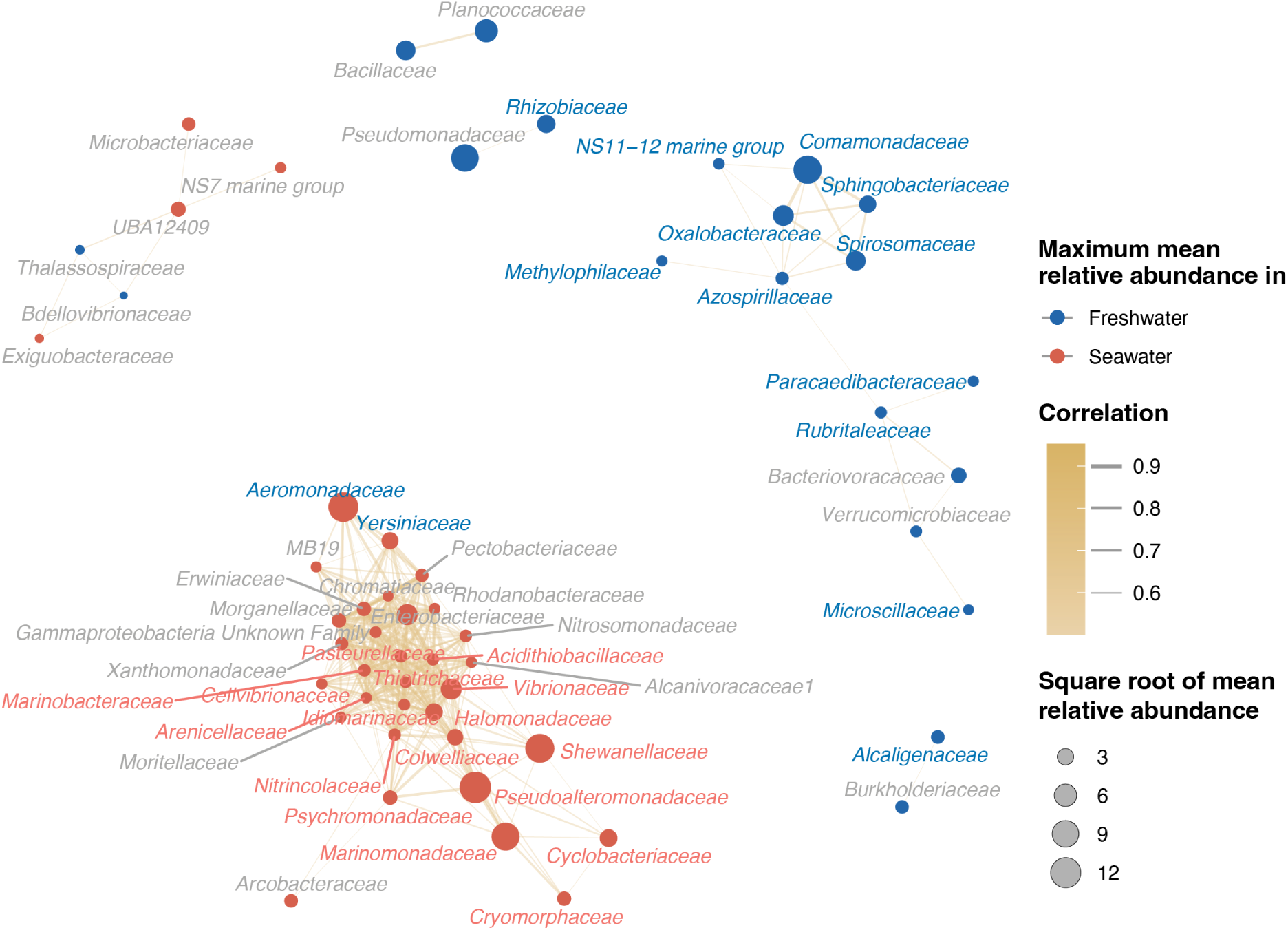
Co-occurrence network of bacterial families across all samples. The network was generated using [SpeSpeNet](https://utrecht-university.shinyapps.io/SpeSpeNet_v1/) and visualized with a Fruchterman–Reingold layout. Nodes represent bacterial families; edges indicate significant correlations (Spearman’s ρ > 0.5). Node colors indicate environmental preference based on the maximum mean relative abundance across all samples. Names in blue and red indicate families enriched in the freshwater or seawater source communities, respectively, which is based on ALDEx2 analysis. Edge width and color correspond to correlation strength (Spearman’s ρ).

The emergence of modular co-abundance patterns is consistent with observations from mixed microbial communities, where functionally or ecologically compatible groups form tightly linked modules (Sierocinski et al. 2017). This is in line with the conceptual hypothesis that modular structure originates from source communities, rather than the newly formed coalesced community (Castledine et al. 2020). Studies on synthetic bacterial communities showed that certain bacterial invaders are more successful when introduced alongside other compatible microbes. These co-introduced microbes can facilitate each other’s establishment through cross-feeding or by altering the local environment to favor their mutual survival, especially in communities with low initial diversity (Dooley et al. 2024). Such synergistic interactions among co-introduced microbes can enhance invasion success (Rivett et al. 2018), while strongly interacting resident species can instead prevent colonization by invaders (Hu et al. 2025). These findings align with theoretical predictions that cooperative interactions can be advantageous during coalescence, while competitive interactions may hinder survival (Lechón-Alonso et al. 2021). In summary, seawater families appear to form more cohesive and modular networks than their freshwater counterparts, likely reflecting co-selection or stronger potential for taxon interactions. This tight modular structure in seawater is consistent with our observation of stronger mixing ratio dependence in seawater.

### Family response varies with mixing ratio and temporal dynamics

Building on these findings, we further analysed the dynamics of freshwater and seawater families across different mixing ratios, culturing conditions, and over time. Consistent with environmental selection shaping overall community structure, freshwater families (e.g., *Comamonadaceae*, *Moraxellaceae*, and *Alteromonadaceae*) dominated in freshwater environments, while seawater families (e.g., *Marinomonadaceae* and *Pseudoalteromonadaceae*) dominated in seawater environments (Figure 6A). This supports the idea that native taxa in a given environment tend to outcompete introduced ones (van Elsas et al. 2012).

**Figure 6.**
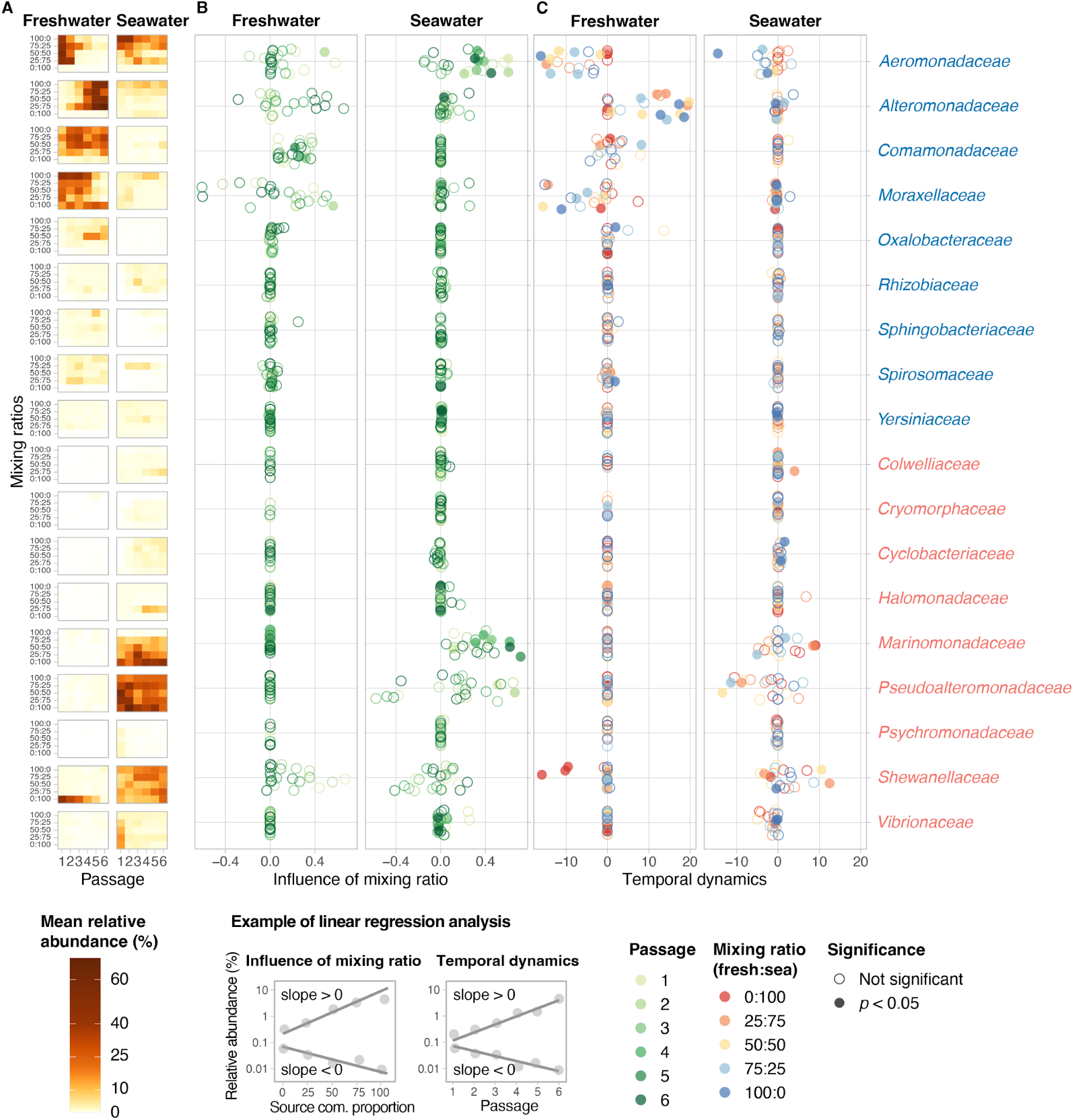
Dynamics in freshwater and seawater families across mixing ratios and over time. Families identified as freshwater- and seawater were determined using ALDEx2 differential abundance analysis. Only families with relative abundance > 1% in at least one sample are shown. (A) Heatmap showing the relative abundance of each family across different passages and mixing ratios (mean of n = 4 replicates per treatment). (B) Regression slopes from linear models of the relationship between relative abundance and the mixing ratio. Specifically, we correlated the proportion of freshwater communities in the mixture for freshwater families, and the proportion of seawater communities for seawater families, see example below. Points are colored by passage number. (C) Regression slopes from linear models of the change in relative abundance over time (passages 1 to 6), see example below. In panel C, points are colored by mixing ratio. Solid circles indicate the significance of the linear regression (*p* < 0.05), while open circles indicate non-significant results. Families are grouped by environmental preference: freshwater families are shown in blue, and seawater families in red.

Most of the freshwater and seawater families showed little change across mixing ratios or over time; only a few showed clear responses to mixing ratios and temporal dynamics (Figure 6, Supplementary Figure S10, S11). This pattern mirrors previous observations that most taxa vary non-monotonically across mixing ratios during coalescence, with only a small fraction responding to mixing ratios in a dose-dependent manner, and other small fractions acting as strong or weak colonisers (Goldman et al. 2025).

In our study, freshwater families such as *Comamonadaceae*, *Aeromonadaceae*, and *Moraxellaceae*, as well as seawater families including *Marinomonadaceae* and *Pseudoalteromonadaceae*, increased in abundance proportionally to the percentage of their native source community in their native environment (Figure 6B). This suggests that their success may depend more on the initial population size than on their capacity for growth under our experimental conditions. As we mentioned above, such inoculation dependence has been proposed to arise from metabolic competition for shared resources between introduced and native microbes (Goldman et al. 2025).

In terms of temporal dynamics, only the freshwater family *Alteromonadaceae* significantly increased over time in its native environment, indicating its slow adaptation and the potential for reestablishment (Figure 6C). *Aeromonadaceae* and *Moraxellaceae* were initially abundant but decreased over time. For seawater families, *Marinomonadaceae* and *Shewanellaceae* showed inconsistent temporal patterns in their native environment, while *Pseudoalteromonadaceae* gradually decreased at lower source community ratios.

Overall, habitat-specific families followed predictable enrichment patterns driven by environmental preference, with a few families being influenced by initial population size or differing by adaptation over time. However, most families were unaffected by these factors, underscoring the complexity of microbial coalescence dynamics, shaped by interactions among taxa, niche breadth, and selection pressure.

## Conclusions

Our mixing experiment demonstrates that environmental filtering is the primary force structuring coalesced freshwater-seawater communities. Mixed communities that grew in freshwater shifted toward a freshwater-like community, and those grown in seawater became more seawater-like. There was asymmetric resilience during coalescence: freshwater communities exhibited greater flexibility in reestablishment, whereas seawater communities depended more on the initial proportion of the inoculated community. The initial proportion of a source community positively impacted the resulting communities only in the native environment. Network analysis revealed a tightly connected module of seawater families, suggesting that they experienced stronger habitat selection or cooperative interactions, while freshwater families were more loosely connected. Taxa responded differently to the initial mixing ratio and time. Few families changed related to the initial mixing ratio. For example, freshwater *Comamonadaceae* and seawater *Marinomonadaceae* and *Pseudoalteromonadaceae* thrived in their native habitat, and their abundance in the coalesced community related to the initial proportion of their source community. Together, these findings highlight the interplay between environmental selection and source community composition in determining coalescence outcomes. Understanding these dynamics is crucial for predicting microbial response to natural mixing events, and for applications in ecosystem management, bioremediation, and microbiome engineering. While our work focused exclusively on bacterial communities, excluding other trophic interactions, such as protists and bacteriophages, which can influence bacterial community dynamics through predation or lysis. Future studies integrating multi-trophic interactions and long-term observations across diverse ecosystems will further refine the general ecological rules governing microbial coalescence and its implications for ecosystem functioning.

## Acknowledgments

We thank Sybille Huck, Clara Nietz, Conor Glackin for their assistance in the field sampling and the rest of the OTC Genomics sampling team for making the sample collection possible; Sandra Studenik for the help in sample collection during the microcosm experiment and test spike-in approach before the experiment; Mia Bengtsson, Martina Herrmann, Martin Taubert, Olga Maria Pérez Carrascal, Kirsten Küsel, Christian Jogler for their comments on the experiment design; Pilleriin Peets for measuring the DOC data; and José Luis López Arcondo for sharing the code to extract bacterial minimal doubling time. ChatGPT was used to assist with writing code for data analysis, and Grammarly was used to check grammar throughout the text.

## Data and code availability

Raw sequencing data generated and analysed by this study are available at the European Nucleotide Archive (ENA) with the study accession number PRJEB94026. All other data and code are available at https://github.com/Jia-Xiu/coalescence_project.

## Contributions

X.J., B.E.D., and T.S. (Schubert) conceived the study. T.S. (Sperlea) and M.L. designed the sampling sites and assisted and provided laboratory capacity during field sampling. X.J. performed the experiments and sequencing with the assistance of T.S. (Schubert), and processed Nanopore raw data with the assistance of S.D. R.B. developed the demultiplexing scripts. A.L. contributed to experimental design and data analysis. P.H. wrote scripts for checking taxa-environment associations in MGnify. X.J. analyzed data, interpreted the results, and drafted the manuscript with input from B.E.D. All authors read, commented on, and approved the final manuscript.

## Supplementary Figures

**Figure S1.**
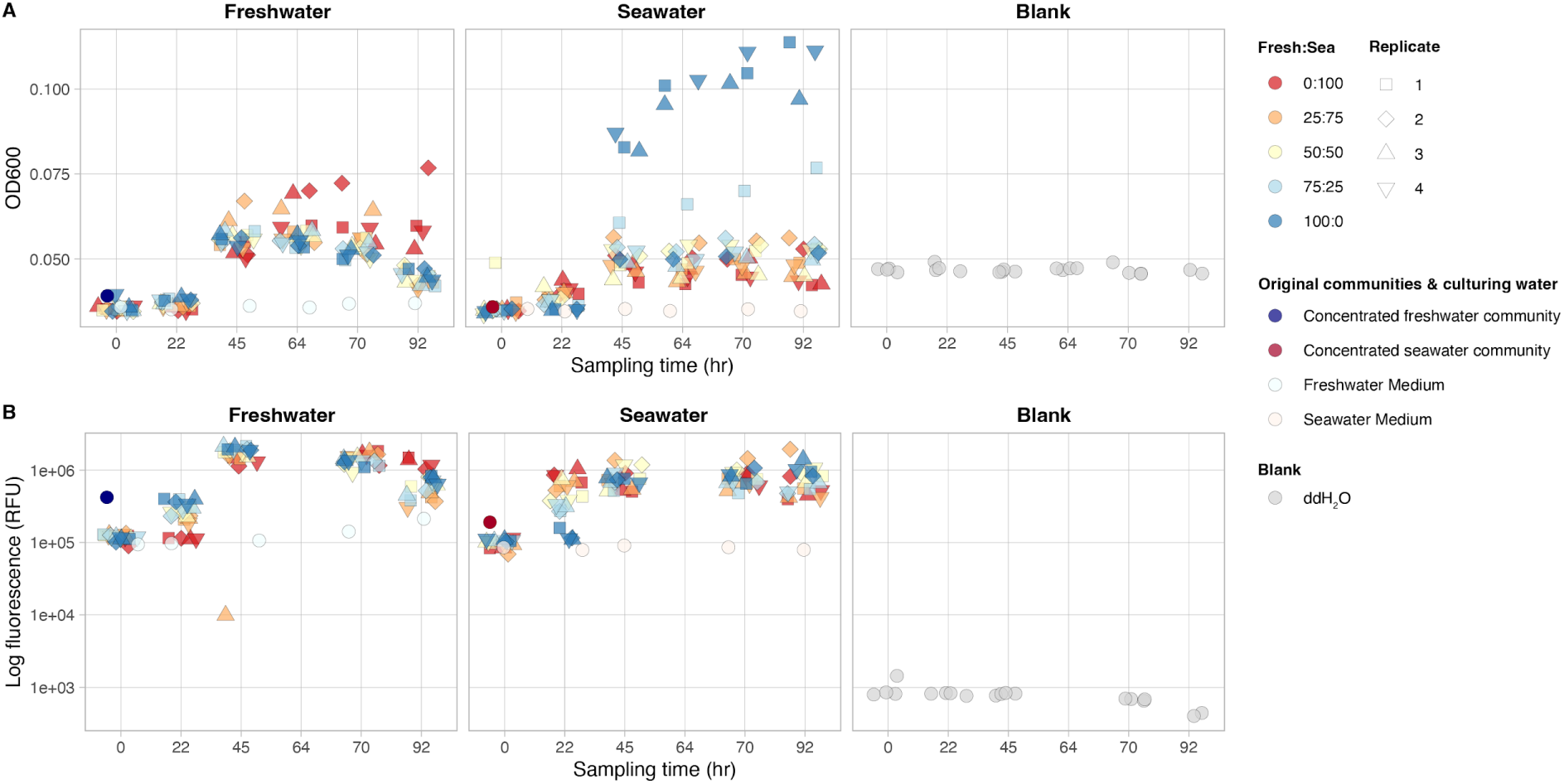
**Cell density dynamics during passage 1**. To track the cell density of the community over time, we measured the optical density (OD 600) and fluorescent signal by SYBR Green dye of all the microcosms at passage 1. Colors represent the mixing ratios of freshwater and seawater source communities, culturing water types, and ddH2O blanks used for density measurements. Shapes represent experimental replicates. The left and middle columns correspond to different culturing water media.

**Figure S2.**
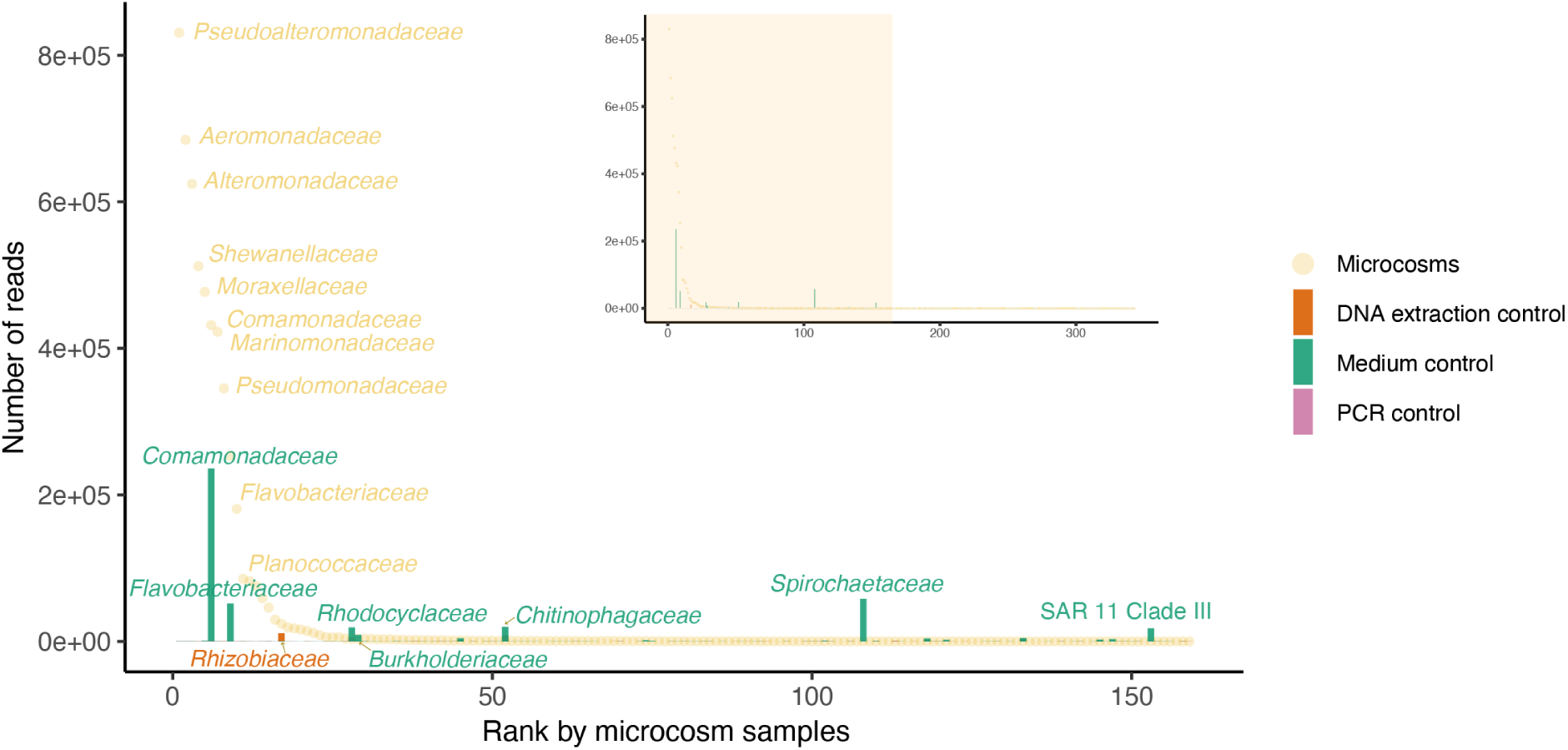
Total raw read counts in microcosm samples and negative controls. The rank abundance curve of bacterial families in all microcosm samples (n = 240) is shown as yellow dots. Corresponding family abundances in negative controls are shown in different colors: DNA extraction controls (orange, n=3), culturing medium controls (green, n = 12; two water media sampled across six passages), and PCR controls (pink, n = 3). The top ten families were labeled on the rank abundance curve. Annotated bars indicating these families in the corresponding control samples had more than 5,000 reads. Our PCR amplification yielded a negligible number of reads compared to the biological sample. DNA extraction controls only had a high detection of Rhizobiaceae. Medium controls still contained detectable bacterial families, indicating that not 100% of microbes from the media were removed. The families in culturing media mostly had fewer reads than in the experimental samples, except for a few families.

**Figure S3.**
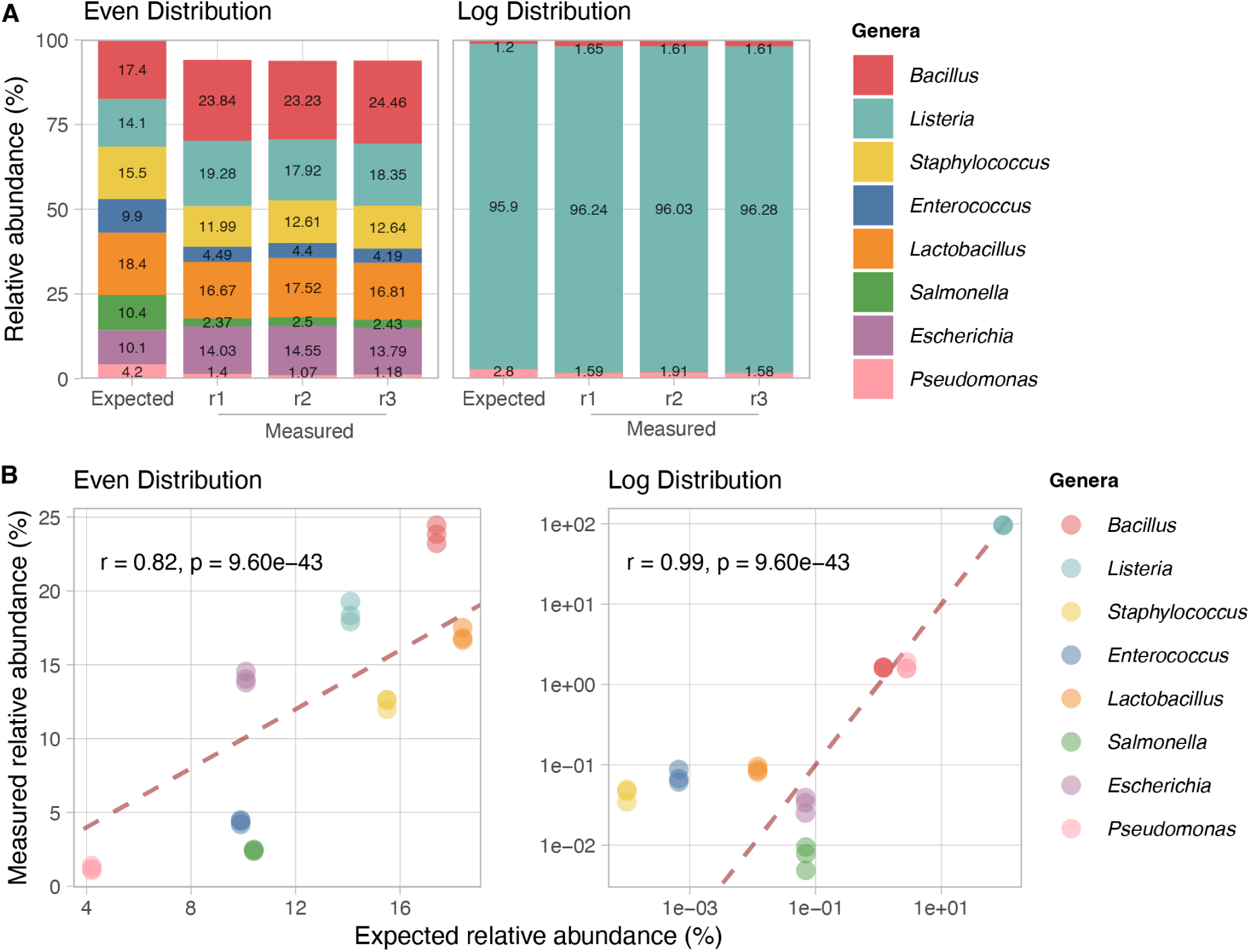
Positive controls of Nanopore sequencing. Comparison of the relative abundance between the expected distribution of the ZymoBIOMICS™ microbial community standard and measured Nanopore sequencing results of the standard at the genus level. (A) Stacked-bar plots and (B) scatter plots show the microbial community standard with even and log abundance distribution. Each real sequencing result is represented by three replicates (r1-r3). Dashed lines in panel B indicate the perfect match between the measured and the expected distribution.

**Figure S4.**
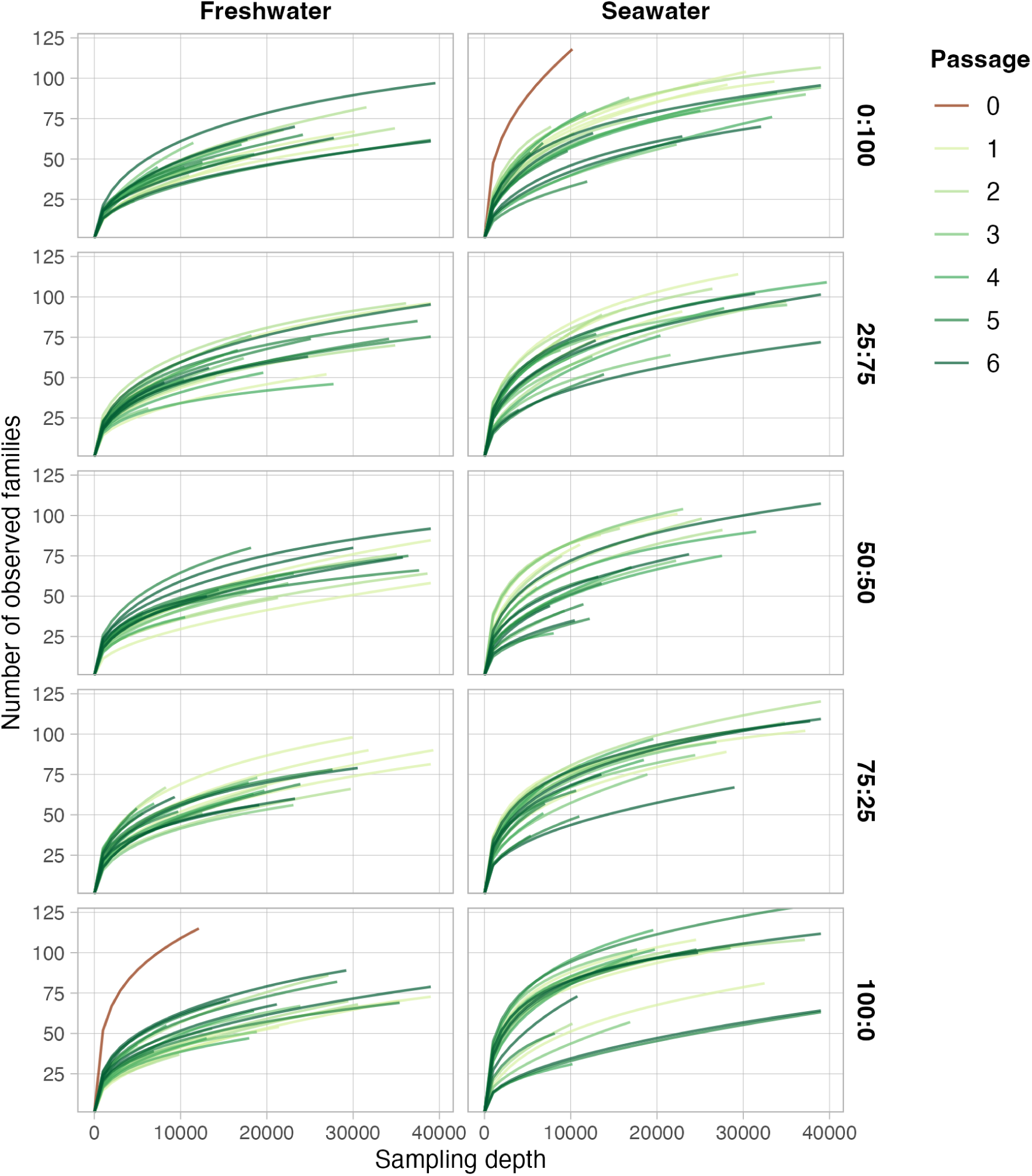
Rarefaction curves of microcosm and field samples. The x-axis shows sampling depth per sample, and the y-axis shows the corresponding number of detected families. Facet rows represent the initial inoculum proportions of freshwater and seawater bacterial communities, while columns indicate the cultural medium of the mixed communities (freshwater or seawater). Colors represent passages, with passage 0 corresponding to the pre-mixing field samples.

**Figure S5.**
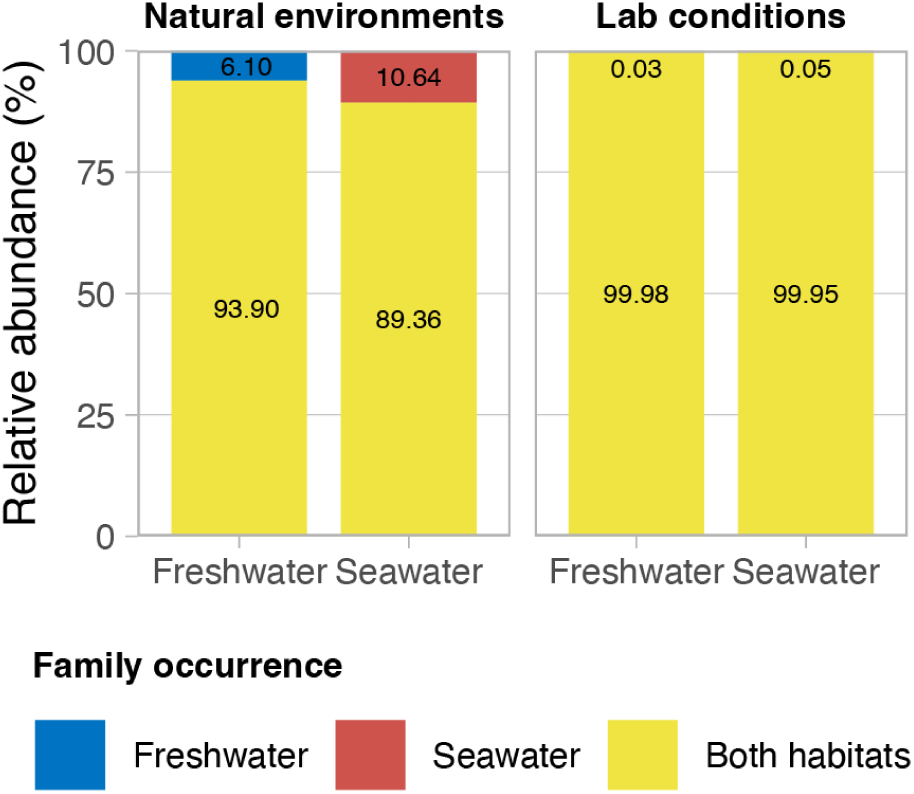
**Total abundance of bacterial families detected in both freshwater and seawater, or exclusive to one habitat under natural or laboratory conditions.**

**Figure S6.**
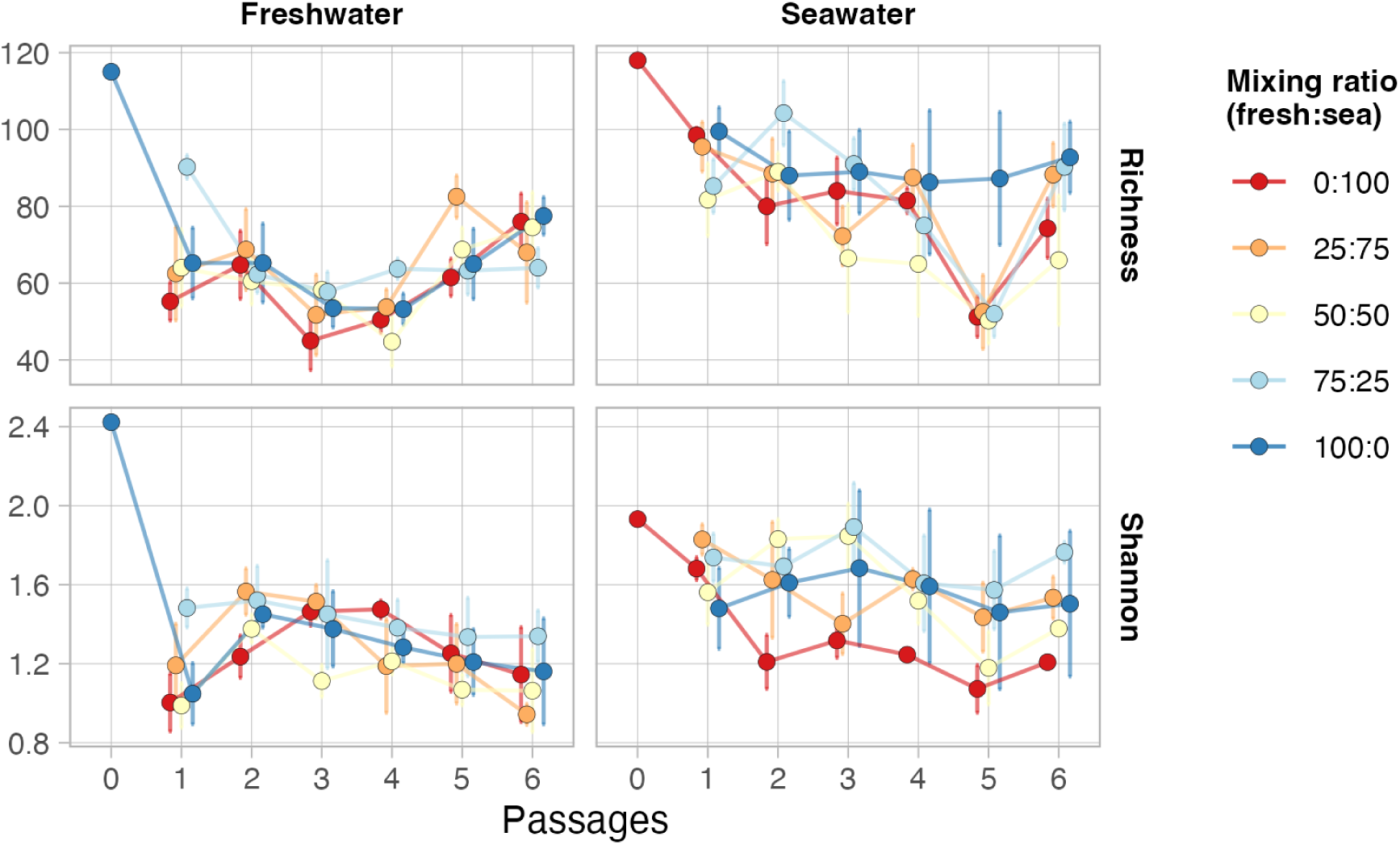
Richness and Shannon diversity of bacterial communities at the family level. Colors represent the mixing ratios of freshwater and seawater communities.

**Figure S7.**
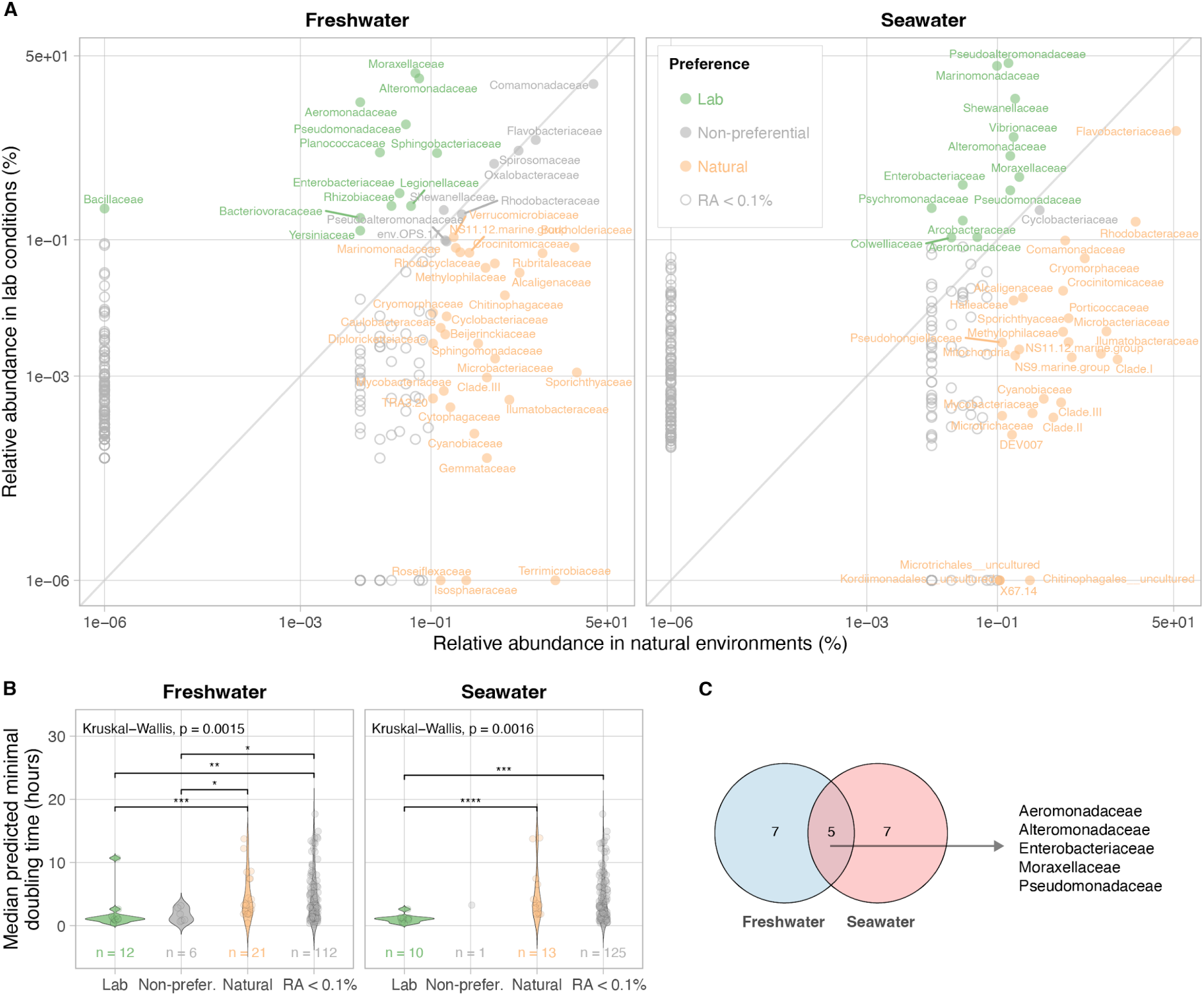
Natural- and lab families in freshwater and seawater source communities. (A) Correlation of relative abundance (RA) between natural and laboratory conditions. Bacterial families were classified as natural- or lab if their log_2_ fold change between these two conditions is greater than 1 or less than −1; families without 2-fold changes were classified as non-preferential. Families with zero abundance were assigned a small pseudocount (10^-6^) to enable the log_2_ transformation. Colors indicate preference groups, and open circles indicate families with RA < 0.1% in both conditions. (B) Predicted growth rates of bacterial families grouped by their preference, based on log_2_ fold changes. (C) Overlap of lab-enriched families identified in both freshwater and seawater environments.

**Figure S8.**
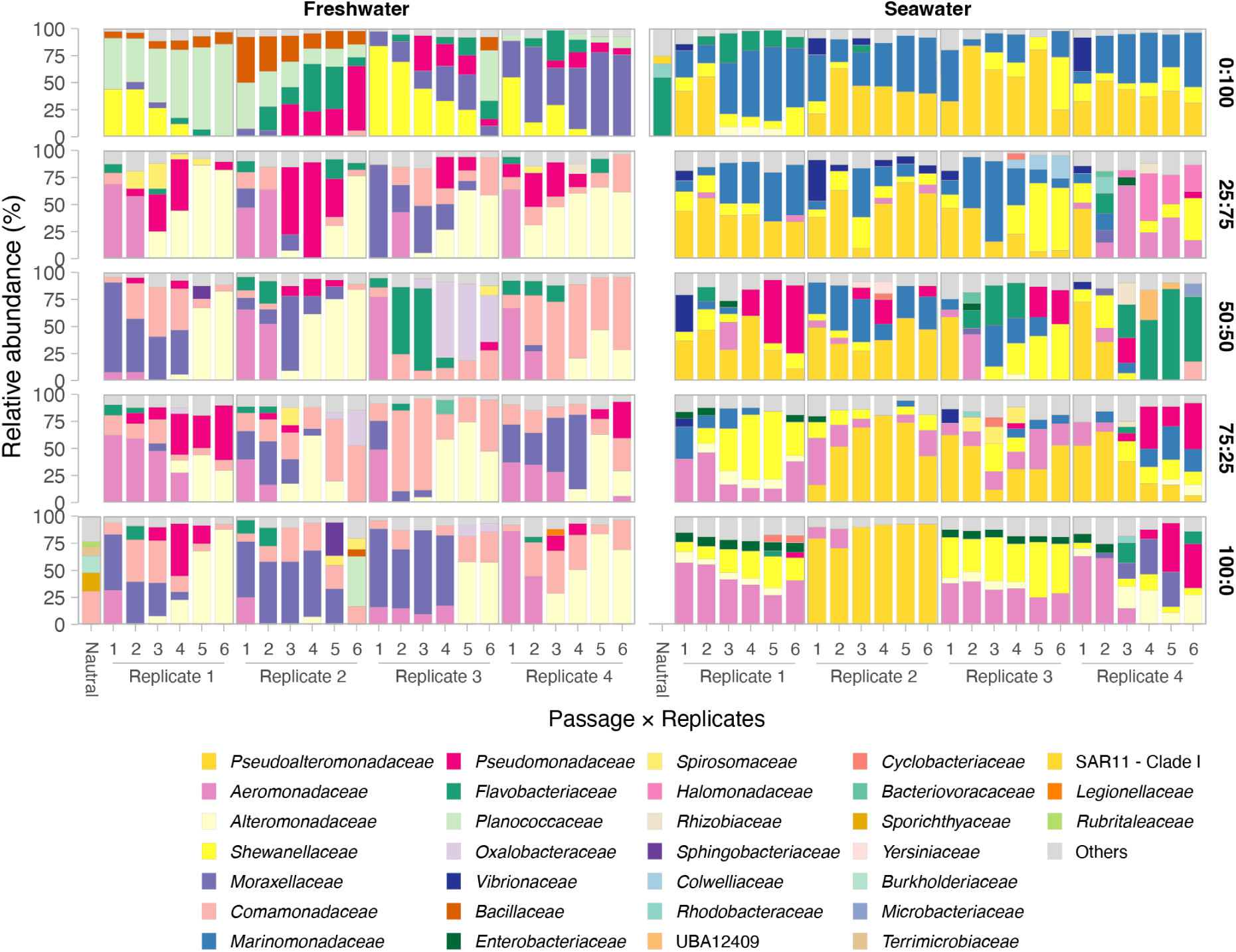
Stacked barplot showing Family composition in mixed communities with relative abundances > 5%. Bar heights represent the relative abundance of each family in individual samples. The x-axis shows the passage number of each replicate. Rows correspond to the initial inoculum proportions of freshwater and seawater bacterial communities. Columns indicate the cultural medium of the mixed communities, either freshwater or seawater.

**Figure S9.**
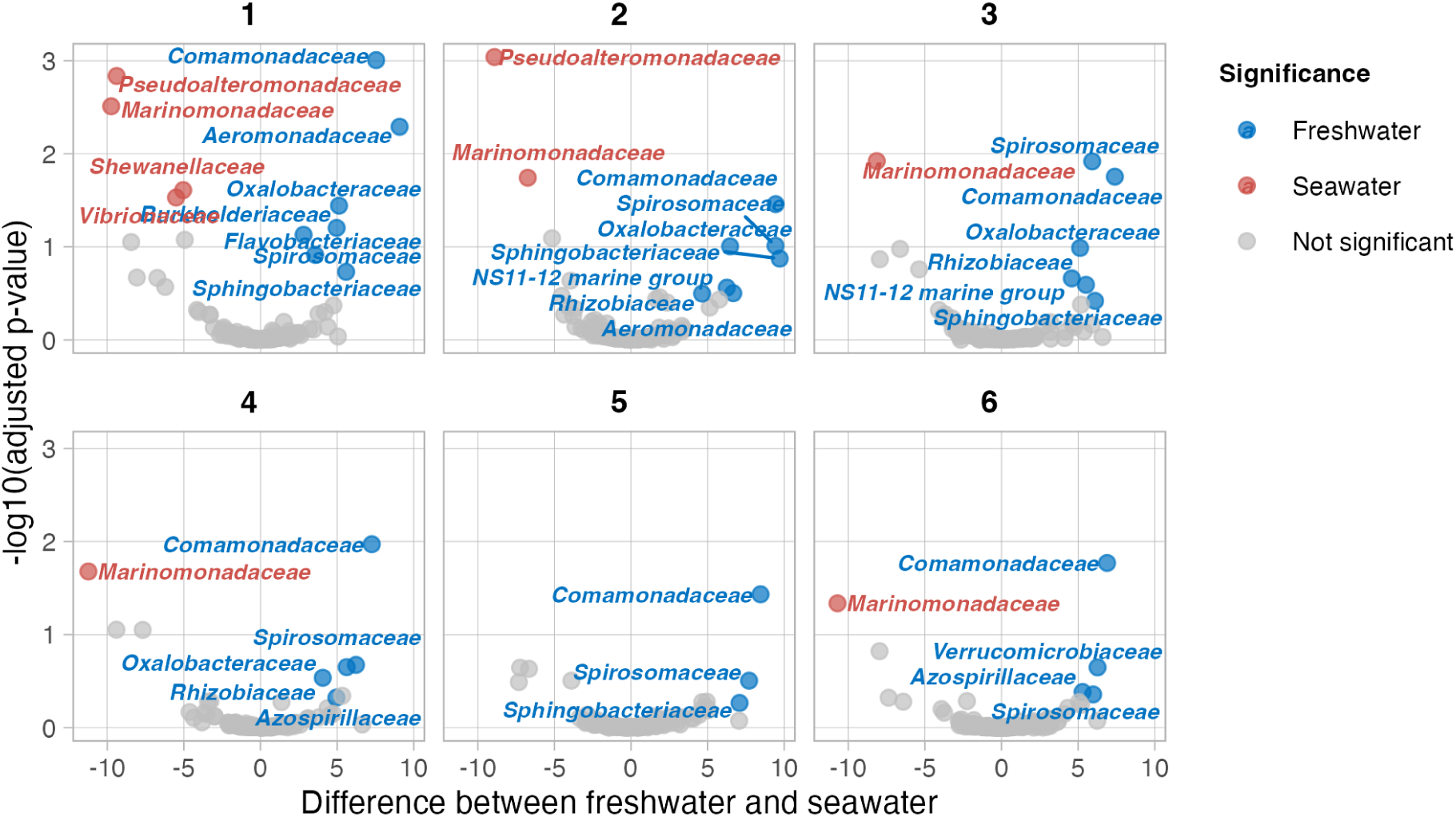
Volcano plot showing differential abundance of families between freshwater and seawater communities across passages under laboratory conditions. Results were based on ALDEx2 analysis, comparing all the passages and replicates between the two different communities under laboratory conditions. The x-axis represents the adjusted *p*-value from Welch’s t-test, and the y-axis shows the median difference in CLR-transformed abundance between freshwater and seawater. Colored dots indicate families with significant differences in either freshwater (blue) or seawater (red), as determined by either the Wilcoxon rank-sum test or the Welch’s t-test (adjusted *p* < 0.05). Each panel corresponds to a different passage.

**Figure S10.** Temporal dynamics of freshwater bacterial families. The x-axis shows the passage number, and the y-axis shows the relative abundance (log_10_-transformed). Each row corresponds to a bacterial family, and each column represents a mixing ratio treatment (e.g., F25:S75 indicates 25% freshwater and 75% seawater community origin). A LOESS curve was fitted to capture temporal trends within each group. Colors indicate the culturing water type (freshwater or seawater), and point shapes present biological replicates.

**Figure S11.** Temporal dynamics of seawater bacterial families. The x-axis shows the passage number, and the y-axis shows the relative abundance (log_10_-transformed). Each row corresponds to a bacterial family; for brevity, “*Pseudoalteromona.*” refers to *Pseudoalteromonadaceae*, and “*Endozoicomona.*” refers to *Endozoicomonadaceae*. Each column represents a mixing ratio treatment (e.g., F25:S75 indicates 25% freshwater and 75% seawater community origin). A LOESS curve was fitted to capture temporal trends within each group. Colors indicate the culturing water type (freshwater or seawater), and point shapes present biological replicates.

## Supplementary Tables

**Table S1.** Location and physicochemical characteristics of the two distinct water types used in this study.

| Original water sample | Freshwater | Seawater |
| --- | --- | --- |
| Latitude | 54°5'2" N | 54°10'55" N |
| Longitude | 12°9'6" E | 12°4'46" E |
| pH | 8.22 | 7.85 |
| Temperature (°C) | 10.7 | 6.7 |
| Conductivity (mS/cm) | 0.60 | 15.23 |
| Salinity (Practical Salinity Unit) | 0.60 | 15.22 |
| Total organic carbon (mg/L) | 11.10 | 4.24 |

**Table S2.** Permutational Multivariate Analysis of Variance (PERMANOVA) results based on Aitchison distance of all bacterial communities in the microcosm experiment. Each treatment factor was identified as a fixed component to the overall model; *p*-values were obtained using 999 permutations; *R*^2^ gives the estimated sizes of components of variation.

| Factor | df | Sum Of Sqs | $R^2$ | Pseudo F | *p* value |
| --- | --- | --- | --- | --- | --- |
| Culturing water | 1 | 8633 | 19.90 | 76.45 | <0.001 |
| Mixing ratio | 4 | 3583 | 8.26 | 7.93 | <0.001 |
| Passage | 5 | 2141 | 4.94 | 3.79 | <0.001 |
| Mixing ratio x Culturing water | 4 | 2803 | 6.46 | 6.21 | <0.001 |
| Mixing ratio x Passage | 20 | 2094 | 4.83 | 0.93 | 0.792 |
| Culturing water x Passage | 5 | 1465 | 3.38 | 2.59 | 0.001 |
| Mixing ratio x Culturing water x Passage | 20 | 2333 | 5.38 | 1.03 | 0.327 |
| Residual | 180 | 20325 | 46.86 |  |  |
| Total | 239 | 43378 | 100 |  |  |

